# Aerobic glycolysis is important for zebrafish larval wound closure and tail regeneration

**DOI:** 10.1101/2021.04.23.441208

**Authors:** Claire A. Scott, Tom J. Carney, Enrique Amaya

## Abstract

The underlying mechanisms of appendage regeneration remain largely unknown and uncovering these mechanisms in capable organisms has far-reaching implications for potential treatments in humans. Recent studies implicate a requirement for metabolic reprogramming reminiscent of the Warburg effect during successful appendage and organ regeneration. As changes are thus predicted to be highly dynamic, methods permitting direct, real-time visualization of metabolites at the tissue and organismal level, would offer a significant advance in defining the influence of metabolism on regeneration and healing. We sought to examine whether glycolytic activity was altered during larval fin regeneration, utilising the genetically encoded biosensor, Laconic, enabling the spatiotemporal assessment of lactate levels in living zebrafish. We present evidence for a rapid increase in lactate levels within minutes following injury, with a role of aerobic glycolysis in actomyosin contraction and wound closure. We also find a second wave of lactate production, associated with overall larval tail regeneration. Chemical inhibition of glycolysis attenuates both contraction of the wound and regrowth of tissue following tail amputation, suggesting aerobic glycolysis is necessary at two distinct stages of regeneration.

## 1 BACKGROUND

While some organisms have the ability to heal scarlessly and regenerate fully functional tissues as adults, others possess this ability only in early developmental stages. Understanding the underlying cellular and molecular processes responsible for successful regeneration may provide essential clues for the development of novel clinical therapies that will promote a better healing and regenerative outcome in humans.

Accumulating evidence indicates metabolism influences complex tissue and cellular processes, including cell differentiation and cell behaviour, however the role of cell metabolism on regeneration has been largely unexplored. Metabolites, such as lactate, have been reported to act as second messengers in cell signalling (Chubanov & Gudermann, 2020), and a switch from oxidative phosphorylation (OXPHOS) to glycolysis is involved in epithelial to mesenchymal transitions (EMTs), which are important for blastema formation in gecko limb regeneration (Gilbert et al., 2013) and in cancer metastasis (Lambert et al., 2017; reviewed in Kalluri & Weinberg, 2009). Work in C. elegans has shown that reduction of mitochondrial activity has positive effects on ageing (reviewed in Y. Wang & Hekimi, 2015), and multiple studies have linked a switch to glycolytic metabolism to the proliferative potential of stem cells (Folmes et al., 2011; Kondoh et al., 2007; reviewed in Mathieu & Ruohola-Baker, 2017). Metabolism further plays an important part in cell identity and differentiation in a variety of settings, including immune cells and neurons (Buck et al., 2016; Zheng et al., 2016). Thus, metabolism plays a wider role in physiology than simply energy production. Given that EMT, proliferation, and differentiation are all processes important for regeneration and wound healing, investigating the potential roles for metabolic reprogramming during regeneration and how these are regulated, may provide insight into how cellular metabolism could be hijacked to facilitate regeneration in humans.

The Warburg effect describes the phenomenon of aerobic glycolysis, in which cells preferentially up-regulate processing of glucose through the conventionally anaerobic pathway of glycolysis and fermentation while decreasing their mitochondrial activity, regardless of the lower energy yield and availability of oxygen (Warburg, 1925). This strategy was originally discovered in cancer cells, but has since been implicated in multiple highly proliferative systems, putatively allowing glycolytic and pentose phosphate pathway (PPP) intermediates to support macromolecule synthesis for new cells (reviewed in Lunt & Vander Heiden, 2011). Since regeneration is highly dependent on cell proliferation and growth, one might expect regenerating cells to employ the Warburg effect to provide for the requirements of forming the new tissues of the regenerate. This appears to be the case in multiple regeneration models. The gene profiles of regenerating Xenopus tails and adult zebrafish hearts show an up-regulation of glycolytic genes and a corresponding down-regulation of mitochondrial genes (Love et al., 2014; Honkoop et al., 2019). This switch to glycolysis has been linked to cell proliferation during cardiomyocyte regeneration (Honkoop et al., 2019).

While varied transcriptomic analyses have suggested that metabolic reprogramming plays a critical role during tissue and appendage regeneration, there is a critical need for improved methods that will facilitate the direct assessment of Warburg-like metabolism during regeneration with temporal and spatial resolution. Recent developments in genetically encoded sensors for various metabolites has advanced the field of metabolic research and has great potential for exploitation in the zebrafish, due to its impressive regenerative capacity (reviewed in Poss et al., 2002 and Poss et al., 2003), combined with the transparency of the embryos. Here we aimed to test the potential of a genetic ratiometric Förster resonance energy transfer (FRET)-based genetic sensor, named Laconic, which is responsive to varying lactate levels (San Martín et al., 2013). Given that rising lactate levels can be used as a measure of aerobic glycolysis / Warburg-like metabolism, we aimed to determine whether this sensor could be used to assess metabolic reprogramming during two models of larval fin regeneration, namely after fin fold and tail amputations. We also aimed to ask whether altering metabolic reprogramming, using chemical inhibitors targeting glycolysis or lactate dehydrogenase, affected the speed or efficiency of wound closure and/or fin and tail regeneration.

## 2 MATERIALS & METHODS

### 2.1 Cloning

Laconic/pcDNA3.1(-) was a gift from Luis Felipe Barros (Addgene plasmid #44238; http://n2t.net/addgene:44238; RRID:Addgene_44238) (San Martín et al., 2013). The Laconic genetic sensor in Laconic/pcDNA3.1(-) was cloned into the pCS2+ vector and the p3 vector from the pTransgenesis system (Love et al., 2011), using standard restriction digest and sticky end recombination methods. In some cases, complimentary primers were designed and annealed to produce short sticky end fragments that were inserted into constructs in order to generate additional complementary restriction sites. For specific restriction enzymes, inserts, and buffers used, see Table S1.

For the transgene cassettes, the modular cloning system pTransgenesis (Love et al., 2011) based on the Gateway system of cloning (Hartley et al., 2000) was used, and recombination was facilitated with the Gateway LR Clonase II Enzyme Mix (Invitrogen, 11791) according to manufacturer instructions: incubated overnight with the LR clonase enzyme at 23°C, followed by inactivation by addition of 0.5µL Proteinase K at 37°C for 10 minutes. 3µL of the reaction was transformed into 30-50µL chemically competent DH5α E. coli cells (Invitrogen) as detailed previously.

### 2.2 Zebrafish husbandry

Adult AB strain wild type and *Tg[ubb:laconic]*^*lkc1*^ zebrafish (Danio rerio) were maintained at 28 °C with a 14 hour light/10 hour dark cycle. Embryos collected from in-crosses were staged as described in Kimmel et al. (1995). All animal experiments were performed in compliance with NACLAR Guidelines of Singapore overseen by the Biological Resource Centre of A*STAR (IACUC Protocol Number 140924), and Home Office guidelines UK. In all cases, embryos were raised in 1X E3 embryo medium as described in Cold Spring Harbor Protocols, or 1X egg water consisting of 60 µg/ml sea salts (Sigma Aldrich S9883), supplemented with 0.1% Methylene Blue unless stated otherwise.

### 2.3 mRNA microinjections

Wild type strain AB zebrafish embryos were injected at the one cell stage into the cell cytoplasm with 1ng laconic sensor mRNA in nuclease free water with phenol red. Laconic sensor mRNA was synthesised from pCS2 plasmids linearised with NotI (NEB), with mMESSAGE mMACHINE SP6 Transcription Kit (Ambion) and purified with lithium chloride (LiCl) extraction.

### 2.4 Generation of transgenic lines

Wild type strain AB zebrafish embryos were injected at the one cell stage into the cell cytoplasm with 25pg tol2 mRNA and 25pg circular plasmid in 1nL. Tol2 mRNA was synthesised from pT3-Tol2 linearised with SmaI (NEB) with mMESSAGE mMACHINE T3 Transcription Kit (Ambion) and purified with LiCl extraction. Injected embryos with strongest expression of mosaic GFP were grown to adults and out-crossed to screen for germline transmission.

### 2.5 Biochemical lactate assay

A commercially available colorimetric lactate assay kit (MAK064, Sigma Aldrich) was used and protocol adapted for embryonic samples. Lactate in the sample reacts with the enzyme mix provided in the kit, the product of which interacts with the supplied lactate probe to produce colour (A570) and fluorescence (excitation/emission = 535/587nm). I chose to identify lactate concentration by measuring the colorimetric product of the enzymatic reaction with lactate at an absorbance of 570nm.

Samples were prepared by flash freezing on dry ice and macerating 25 dechorionated eggs or embryos with a plastic micropestle in 45µL 2:2:1 acetonitrile:methanol:dH2O at -20°C or pre-chilled on dry ice. Samples were then centrifuged at 4°C at 15000rcf for 10 minutes, the supernatant collected into a new tube and stored at -20°C until use in the assay. 5µL of the embryo supernatant was used per reaction.

A standard curve was set up using known concentrations of a lactate standard (0, 2, 4, 6, 8 and 10nM per reaction) with the addition of 5µL 2:2:1 to each reaction in order to control for any background or change in enzyme activity caused by the buffer.

Triplicate reactions were set up otherwise according to manufacturer instructions, with a minimum of three biological repeats. Reaction incubation time was extended to 3 hours, and absorbance at 570nm (A_570_) was read on a microplate reader (BioTek Synergy H1) in triplicate to give a total of nine readings per sample.

### 2.6 Fin fold and tail amputations

2 days post fertilisation (dpf) embryos were mounted in 1% low melting agarose (Invitrogen 16520100) supplemented with 0.04% MS-222 (tricaine, Sigma Aldrich E10521) on a glass microscope slide for imaging with an upright microscope or in a 35mm glass bottomed dish (Thermo Scientific Nunc) for imaging with an inverted microscope, and imaged pre-amputation. Amputations were made while embryos were mounted in agarose with either a size 10 or 15 scalpel blade, the agarose surrounding the fins excavated, and the embryos covered with media. Fin fold amputations were performed just distal to the tip of the notochord, and tail amputations were oriented using the pigment gap and transected just distal to the circulatory loop of the caudal vein (Fig. S3A,B). Images were then taken at various timepoints post amputation (Fig. S3C,D) with the embryos being de-mounted from the agarose and kept in 1X egg water or 1X E3 at 28°C between imaging timepoints of longer than one hour.

For experiments where imaging immediately post amputation was not required, 2dpf embryos were not mounted in agarose, and instead amputated in a droplet of 1X egg water supplemented with 0.04% MS-222 on a glass microscope slide, transferred to 1X egg water with or without drug treatment within 5 minutes of amputation, and maintained at 28°C.

### 2.7 Microscope sample preparation

Embryos were visually screened using a fluorescent dissecting microscope for transgenic expression and dechorionated manually with forceps. Embryos were mounted in 1% low melting agarose (Invitrogen 16520100) supplemented with 0.04% MS-222 (tricaine, Sigma Aldrich E10521) on a microscope slide for imaging with an upright microscope, or in a 35mm glass bottomed dish (Thermo Scientific Nunc) for imaging with an inverted microscope.

### 2.8 Microscopy

Images were acquired on an AxioImager.M2 upright microscope (Zeiss) for figures 1 to 6 using a 5X/0.16 EC Plan-Neofluar, 10X/0.3 EC Plan-Neofluar, or 20X/0.4 Corr LD Plan-Neofluar objective was used as specified. Zeiss filter sets for CFP (BS455) and FITC/mCherry (DBS525/50 + 650/100) were utilised for Laconic imaging, with Violet (430nm) excitation from a Colibri 7 LED fluorescent light source. Imaging software: Zen Blue 2.3 Pro. The images were collected using a 2.8 Megapixels (Axiocam 503) colour camera at 14-bit on the black-and-white setting at room temperature.

**Figure 1.**
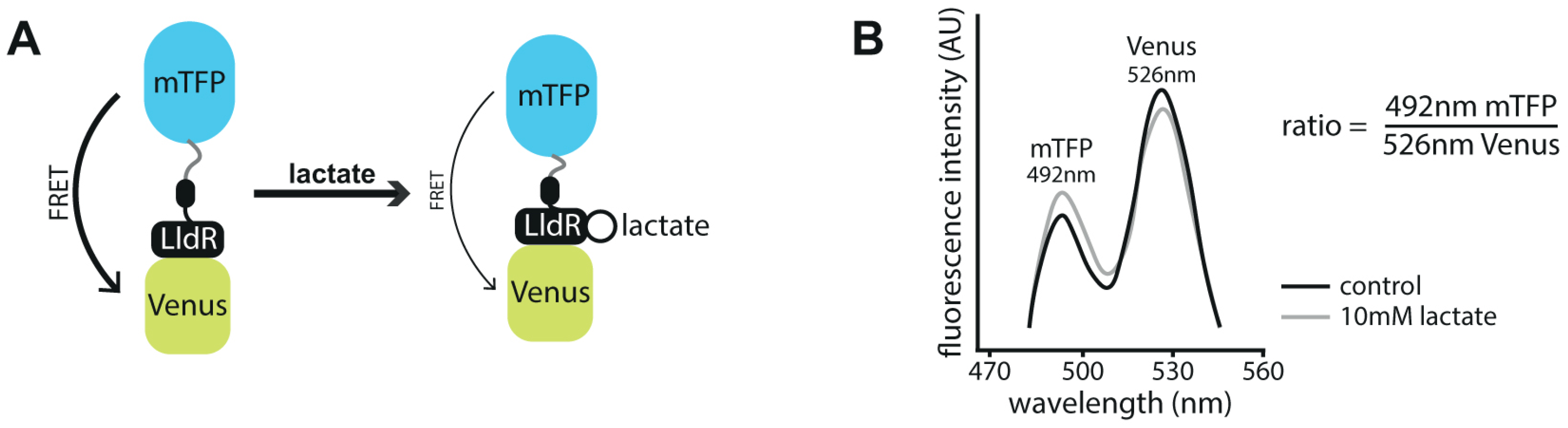
Schematic of the Laconic genetically encoded sensor for lactate. (A) Schematic depiction of the Laconic genetically encoded FRET-based sensor. A conformational change upon lactate binding to the LldR lactate binding region fused to the Venus chromophore induces a change in energy transfer efficiency from mTFP to Venus. (Adapted from San Martín, A. et al., 2013. Fig.1B) (B) Graph depicting emission spectra of Laconic. When bound to lactate, fluorescence intensity detected in the range of mTFP increases and that of Venus decreases due to reduced FRET efficiency, thereby increasing the Laconic ratio as calculated by mTFP/Venus. (Adapted from San Martín, A. et al., 2013).

Laconic imaging for Figure 7 was acquired on an Eclipse Ti inverted microscope (Nikon) with a 4X/0.13 Plan Fluor PhL DL objective using a SpetraX light engine (Lumencore) with individual Semrock emission filters for eCFP (480/30) and eYFP (535/30), and excitation with Blue (440/20) LED fluorescent light source and filter. The images were collected using a Retiga R6 (Q-Imaging) CCD camera at 14-bit. Imaging software: NIS Elements AR.46.00.0. Mechanised point visiting was used to allow multiple positions to be imaged and environment was maintained at 28°C.

Images for immunohistochemistry and phalloidin staining were acquired on a LSM800 (Zeiss) upright confocal microscope using a 20X/0.5 N-Achroplan WD (water) objective. Emission was collected at 400-550nm with excitation laser 488nm for pNM-488, 561-700nm emission with 561nm excitation for rhodamine phalloidin, and 400-454nm emission with 405nm excitation for DAPI. Imaging software: Zen Blue 2.3 Pro. The images were collected using two-channel multi-alkali PMT detectors at 8-bit, pinhole 1AU, with Z-stacks of 2.44µm slice intervals.

### 2.9 Image analysis

All processing of images for calculating ratio and measuring fluorescence or ratio was conducted in Fiji (version 2.0.0). Average background was subtracted and threshold applied to remove remaining background, then Laconic ratio calculated by dividing the 428nm emission channel by the 485nm emission channel using the Image Calculator function. Pseudo-colouring was applied using Lookup Table “16 colors”.

Quantification of fin width or regrowth length for amputation experiments was conducted with Fiji. Regrowth length was defined as the perpendicular distance from the tip of the notochord to the distal fin fold edge.

### 2.10 Pharmacological treatment

2dpf embryos were maintained in 0.5X E2 medium (half strength modification of the E2 embryo medium described in Cunliffe (2003)) in place of 1X E3 embryo medium. Drug treatments were maintained until one hour post amputation.

Antimycin A (AA, Sigma Aldrich A8674) was dissolved to make a stock solution of 5mM in dimethyl sulfoxide (DMSO, Sigma Aldrich D8418) and diluted 1:1000 in E2 medium for a working concentration of 5µM with a final concentration of 0.1% DMSO.

A stock concentration of 500mM Sodium oxamate (Sigma Aldrich O2751) was dissolved in distilled water fresh for each use and diluted in E2 supplemented with 0.04% MS-222 (tricaine, Sigma Aldrich E10521) to a working concentration of 10, 150 or 200mM. For amputation experiments examining the initial hour of regeneration, embryos were amputated in media containing oxamate at a concentration of 150 or 200mM; for those regarding the whole of the regeneration process, embryos were placed into media containing 10mM oxamate immediately following amputation and kept in the drug until assessment at 120hpa.

A stock concentration of 4mM GNE-140 (Sigma Aldrich SML2580) was dissolved in DMSO and diluted in E2 supplemented with 0.04% MS-222 (tricaine, Sigma Aldrich E10521) to a working concentration of 40mM and 400mM. For amputation experiments examining the initial hour of regeneration, embryos were amputated in media containing GNE-140 at a concentration of 400mM; for those regarding the whole of the regeneration process, embryos were placed into media containing 40mM GNE-140 immediately following amputation and kept in the drug until assessment at 120hpa.

For sodium azide (NaN_3_, Sigma Aldrich S2002) treatment, powder form of the drug was dissolved in 1X phosphate buffered saline (PBS, Sigma Aldrich P5493) fresh for each use at a stock concentration of 1.5M and diluted in E2 supplemented with 0.04% MS-222 to a working concentration of 15mM or 25mM.

2-Deoxy-D-glucose (2DG, Sigma Aldrich D8375) was dissolved in distilled water to a stock solution of 250mM and diluted 1:10 in 1X egg water with methylene blue to produce a working concentration of 25mM. In amputation experiments, embryos were placed into media containing 2DG immediately following amputation and 2DG treatment was maintained from 0hpa until 72hpa (Fig. S1C), then washed out and the embryos placed in new 1X egg water with methylene blue, as longer treatment results in embryo mortality. For Laconic imaging experiments of the first hour post amputation, amputations were made while embryos were mounted in agarose with either a size 10 or 15 scalpel blade, then agarose surrounding the fins was excavated and the embryos covered with the treatment solution. Images were then taken 10 minutes post amputation. For experiments over the whole of regeneration, embryos were amputated in a droplet of tricaine solution on a glass microscope slide and transferred to inhibitor treatment within five minutes post amputation. For immunohistochemistry samples, 2dpf embryos were amputated in a droplet of the oxamate and tricaine solution on a glass microscope slide with a size 10 or 15 scalpel blade, incubated for 10 minutes at room temperature, then fixed as described below.

### 2.11 Immunohistochemistry

10 AB strain wild type embryos per condition per experiment were fixed in either 4% paraformaldehyde (PFA, Sigma Aldrich F8775) in 1X phosphate buffered saline (PBS, Sigma Aldrich P5493) at room temperature for 2 hours or 95% methanol (MeOH, Sigma Aldrich 34860)/5% glacial acetic acid (GAA, Sigma Aldrich A6283) at -20°C for 4 hours. In brief: if fixed in 95% MeOH/5% GAA washes were done with PBDT (1XPBS/1%BSA/2%DMSO [dimethyl sulfoxide, Sigma Aldrich D8418]/0.5%Tween, if fixed in 4% PFA washes were in PBST (1XPBS/0.1%Tween or Triton). Samples were systematically rehydrated in methanol washes, then washed in PBDT or PBST as specified by the fixation method, followed by acetone cracking at -20°C for 7 (PFA fix) or 14 (MeOH/GAA fix) minutes. Blocking was in 2% donkey serum for 2 hours at room temperature. Primary antibody, rabbit α-phosphomyosin light chain II (Ser19) (Cao et al., 2017; Araya et al., 2019, Cell Signalling Technology #3671), was added in fresh block at a dilution of 1:250 and incubated overnight at 4°C. Secondary antibody, donkey α-rabbit Alexa Fluor 488 (Invitrogen R37118), was added in fresh block at a dilution of 1:500 and incubated overnight at 4°C. DAPI (Invitrogen D1306) was added 1:1000 (10nM) during the first half an hour wash post-secondary antibody addition. Samples were transferred to 50% glycerol (Sigma Aldrich G5516) in 1X PBS and stored at 4°C.

Phalloidin staining: rhodamine phalloidin (Invitrogen R415) was made up as 40X stock solution in methanol and added in a 1:40 dilution to 4% PFA fixed samples, either alone directly after fixation or in tandem with secondary antibody addition.

### 2.12 Statistical analysis

GraphPad Prism 8 was used for statistical testing, with sample numbers exceeding 6 in all experiments, and each experiment was replicated three or more times. Column or grouped statistics and analyses of differences between means were implemented for all data sets. For column statistics, two-tailed unpaired t-tests with assumed Gaussian distribution were used. Two-way ANOVA was used with Sidak’s multiple comparisons test to compare means between groups. All data are presented as mean±s.d., and differences were considered significant to * at P<0.05, ** at P<0.01, *** at P<0.001, and **** at P<0.0001. Not significant (ns) was considered P≥0.05, 95% confidence interval.

## 3 RESULTS

### 3.1 Laconic can be used to monitor lactate levels in zebrafish embryos and larvae

The genetically encoded biosensor, Laconic, was originally developed and tested on cells in culture (San Martín et al., 2013) and has since been used in mouse brains (Mächler et al., 2016), but has not been used in whole organisms, such as the zebrafish. The FRET-based sensor is composed of a lactate binding region, the bacterial transcription factor LldR, linked with the fluorescent proteins mTFP and Venus (San Martín et al., 2013). Upon binding of lactate, a conformational change decreases the FRET efficiency of energy transfer from the donor chromophore, mTFP, to the acceptor chromophore, Venus (Fig. 1A). By exciting mTFP and measuring the emission from both mTFP and Venus, one can form a ratio to depict the changes in lactate with temporal and spatial resolution. The mTFP/Venus ratio (the Laconic ratio) increases with lactate levels (Fig. 1B).

We devised a positive control, whereby we treated embryos 17 hours post fertilisation (hpf) with antimycin A (AA), a mitochondrial OXPHOS inhibitor which acts to drive glucose into aerobic glycolysis and thus conversion into lactate. We first established the action of AA was indeed increasing lactate levels by measuring the concentration of lactate in treated embryos using a commercial biochemical assay kit. When comparing lactate concentrations in embryos before and after 10 minutes of treatment with either AA or DMSO, we found that AA treated samples gave significantly higher readings of lactate concentrations (Fig. S1A). From this we were confident that AA treatment would provide a satisfactory method to test the efficacy of the Laconic sensor.

In order to confirm Laconic reports lactate dynamics in the zebrafish embryo, we injected in vitro transcribed *laconic* mRNA into one cell stage zebrafish embryos and imaged them before treatment and after one hour of treatment, either with AA or DMSO vehicle control. The Laconic ratio increased significantly in response to AA but not DMSO (Fig. S1B-D). Thus, we concluded that Laconic can successfully report lactate levels in zebrafish embryos.

We then generated a transgenic line (*Tg[ubb:laconic]*^*lkc1*^) expressing Laconic under the control of the *ubiquitin B* promotor (*ubb*) (Mosimann et al., 2011) in order to visualise lactate levels over the course of larval fin regeneration. This Laconic transgenic line showed a higher Laconic FRET ratio following treatment with AA, indicating it reliably reports lactate levels in embryos and larvae (Fig. S2A-C). As further confirmation, we performed a second positive control using an alternate mitochondrial inhibitor, sodium azide (NaN_3_), which acts on complex IV of the electron transport chain (Bennett et al., 1996), similarly blocking OXPHOS, and driving the cell towards anaerobic respiration. As with AA, one hour of treatment with NaN_3_ increased the Laconic ratio significantly compared to PBS control treatment. Furthermore, as NaN_3_ is a reversible inhibitor, we also measured Laconic ratio after 24 hours of recovery following washout of the drug and observed lactate levels had returned to those of controls (Fig. S2D-F).

### 3.2 Lactate levels increase transiently immediately post amputation

We next assessed whether lactate levels change following two types of larval fin injury, namely following distal fin fold amputation, where epidermal tissue was excised, and following tail amputations, where many additional tissues are transected, including the notochord, spinal cord, major blood vessels and skeletal muscle (Fig. S3A,B). Both types of injury induce three similar phases of tissue response and regeneration (wound healing, proliferation, and outgrowth and differentiation, Fig. S3C,D) that are comparable to adult fin regeneration, albeit on a faster time scale (Kawakami et al., 2004; Mateus et al., 2012; Romero et al., 2018), allowing findings made in zebrafish larvae to likely be transferrable to adult systems and other organisms. Upon amputation, most regenerative responses form a highly proliferative structure termed the blastema (reviewed in Londono et al., 2018). Since regeneration involves the regrowth of lost tissue, it would stand to reason that larval tail regeneration would be dependent on the generation of significant amounts of new biomass. As such, appendage regeneration is a good candidate for the Warburg effect (Love et al., 2014). Thus, we aimed to image lactate levels, as a proxy for the Warburg effect, during regeneration following larval zebrafish fin fold and tail amputations utilising the *Tg[ubb:laconic]*^*lkc1*^ line.

We found that following distal fin fold amputation, Laconic ratio increased immediately following amputation, peaking within the first five minutes post amputation (mpa) and returning to control levels by 50mpa (Fig. 2A,C,D). Spatially, lactate levels were raised in a broad gradient from the wound border, up to approximately 80µm into the fin. For the remaining period of distal fin fold regeneration there was no significant difference between amputated and un-amputated controls (Fig. 2B,E). The first event following amputation is the wound healing phase, in which the wound rapidly closes within 10 minutes by a purse-string contraction mediated by an actomyosin cable (Mateus et al., 2012) (Supplementary Movie S1). Glycolysis is able to produce ATP more rapidly than OXPHOS, which is why the fastest contracting muscle fibres, which are also actomyosin based, are largely glycolysis-based in their metabolism (reviewed in Schiaffino & Reggiani, 2011). Potentially this is an explanation for the transient burst of lactate and glycolysis activity after amputation, as a strategy for swiftly producing large quantities of ATP to fuel the contraction of the actomyosin cable during wound closure.

**Figure 2.**
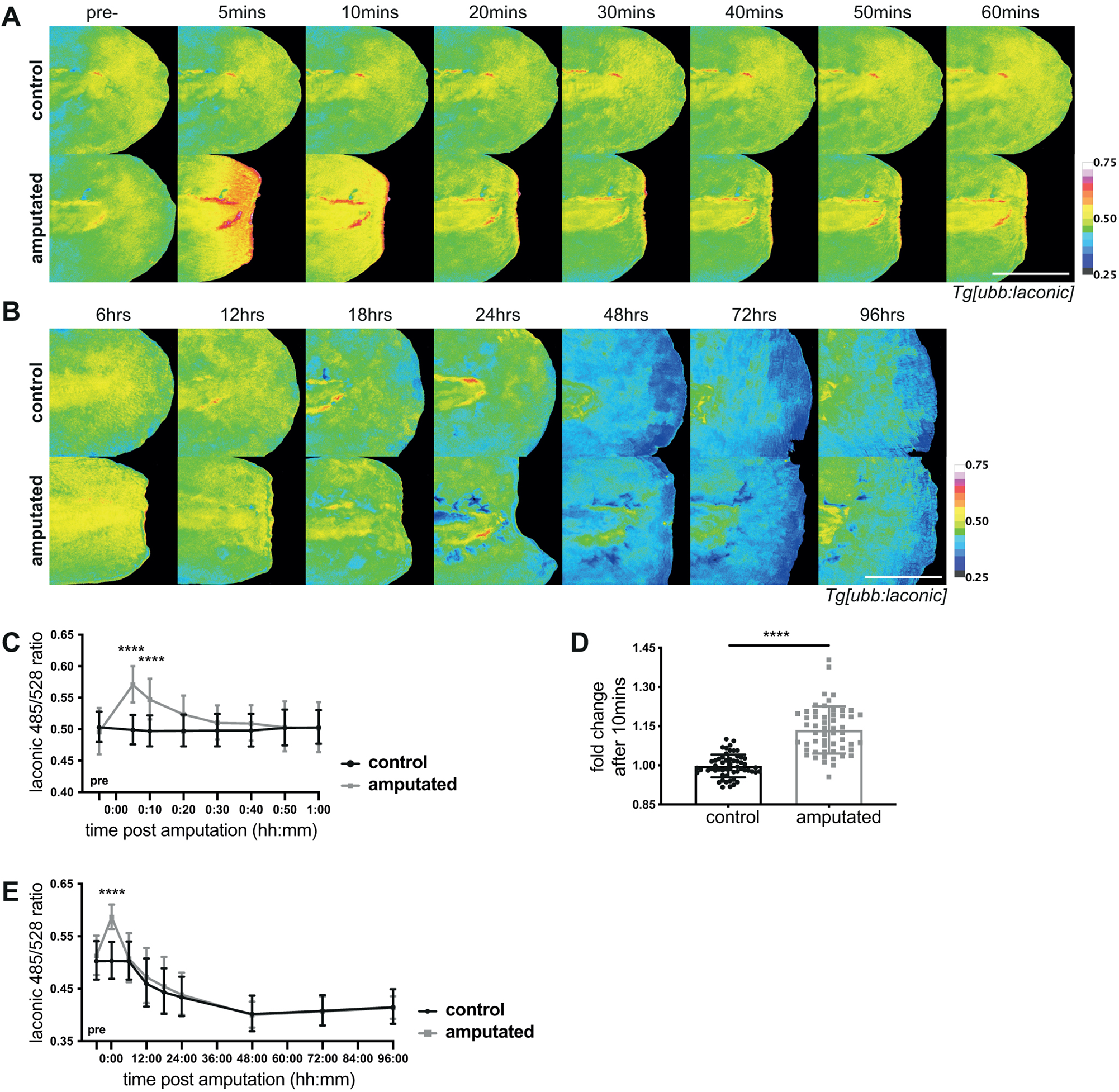
Lactate levels in fin fold regeneration. A) Micrographs of representative *Tg[ubb:laconic]*^*lkc1*^ embryos tails at 48hpf imaged pre-amputation and the same individual embryo followed over the course of one hour post amputation, pseudocoloured to show Laconic ratio. (B) Micrographs of representative transgenic *Tg[ubb:laconic]*^*lkc1*^ embryos tails amputated at 48hpf and imaged at given time-points over the course of regeneration until five days post amputation, pseudocoloured to show Laconic ratio. (C) Graph showing quantification of raw Laconic ratios pre-treatment and over the course of one hour post amputation. Two-way ANOVA to calculate significance, n=18. (D) Graph showing fold change between pre-amputation ratio and 10 minutes post amputation. Students ‘t-test to calculate significance, n=56. (E) Graphs showing quantification of raw Laconic ratios pre-treatment and at various timepoints post amputation. Two-way ANOVA to calculate significance, n=18. All scale bars represent 200μm. Differences were considered significant to * P<0.05, ** P<0.01, *** P<0.001, **** P<0.0001, and ns P≥0.05.

### 3.3 Inhibition of lactate production inhibits wound contraction

We tested this hypothesis by inhibiting aerobic glycolysis and assessing the effects on the actomyosin cable contraction at the wound. To do this, we transiently inhibited the activity of lactate dehydrogenase (LDH) using the competitive inhibitor, sodium oxamate (M. Zhou et al., 2010). LDH converts pyruvate to lactate and regenerates NAD+, permitting continued glycolysis independent of mitochondrial activity. We reasoned that using chemical inhibitors provide a powerful method of transiently inhibiting LDH activity, not easily afforded by alternative methods, such as genetic knockouts or knockdown, which are likely incompatible with survival. To test the efficacy of the drug, we first asked whether oxamate affected the rapid increase in lactate levels following injury. Oxamate treatment indeed prevented the increase in lactate post injury in a dose dependent manner (Fig. 3A,C,E). We then asked if this LDH inhibition affected wound healing / closure or subsequent fin fold regeneration. Indeed, oxamate treatment potently inhibited wound contraction following injury (Supplementary Movie 2). To quantify this effect, we measured fin width across the plane of amputation as a description of wound contraction and found that oxamate treatment, following fin fold amputation, resulted in the wound remaining significantly wider at 10mpa (Fig. 3B,D,E). Removing oxamate from the media after one hour post amputation (hpa) did not affect overall regeneration, and fins appeared similar to controls in terms of Laconic ratio and fin regrowth length (Fig.3A-D). These data suggest that rapid glycolysis activity is required for the rapid contraction of the wound margin post amputation, but the embryos are still able to recover following temporary inhibition of LDH and this is not detrimental to long-term regeneration. This reflects our previous Laconic data, which showed only a transient rise in lactate levels following distal fin fold amputation and elevated lactate levels are not sustained during the fin fold regeneration phase. To confirm our hypothesis that LDH activity and aerobic glycolysis is important for wound contraction, we also utilised an alternate LDH inhibitor, GNE-140 (Pusapati et al., 2016), to transiently inhibit LDH activity during the wound healing following distal fin amputation. GNE-140 also strongly inhibited the rapid contraction of the wound post amputation, comparable to the inhibition seen following oxamate treatment (Fig. S4A,B) (Supplementary Movie 3 and 4).

**Figure 3.**
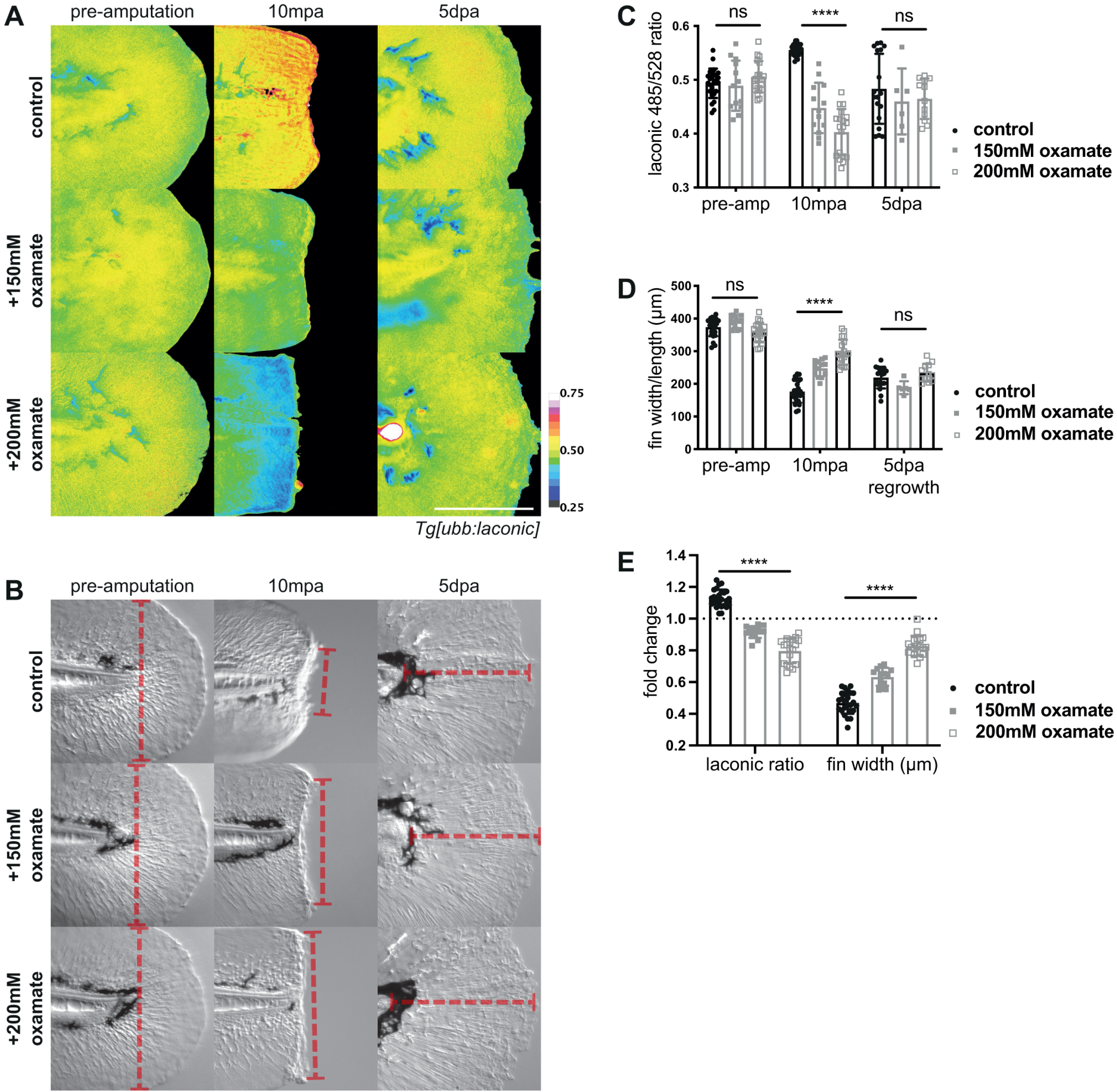
Lactate dehydrogenase inhibition in wound healing. (A) Micrographs of representative *Tg[ubb:laconic]*^*lkc1*^ embryos tails at 48hpf imaged pre-amputation, 10 minutes post amputation with treatment with oxamate or water control, and five days post amputation, pseudocoloured to show Laconic ratio. (B) DIC micrographs of representative transgenic *Tg[ubb:laconic]*^*lkc1*^ embryos tails as in (A) with examples of measurements (red dashed line) taken for fin width and length quantification. Pre-amputation and wound width taken to be from edge to edge of the fin fold just distal to the notochord along the amputation plane; regrowth taken from the end of the notochord to the most distal edge of the fin fold, perpendicular to amputation plane. (C) Graph showing quantification of raw Laconic ratios pre-, 10 minutes post-, and five days post-amputation. Two-way ANOVA to calculate significance, n=25 (control), n=13 (150mM oxamate), n=19 (200mM oxamate). (D) Graph showing measured fin widths/length in micrometres pre-, 10 minutes post-, and five days post-amputation. Fin length measured from the tip of the notochord to the distal edge of the fin fold. Two-way ANOVA to calculate significance, n=25 (control), n=13 (150mM oxamate), n=19 (200mM oxamate). (E) Graph showing fold change (10mpa value divided by pre-amputation value) of Laconic ratio and fin width in micrometres in the first 10 minutes of amputation with treatment with oxamate or water control. Dotted line on the Y axis marks a fold change of 1 (no change). Two-way ANOVA to calculate significance, n=25 (control), n=13 (150mM oxamate), n=19 (200mM oxamate). All scale bars represent 200μm. Differences were considered significant to * P<0.05, ** P<0.01, *** P<0.001, **** P<0.0001, and ns P≥0.05.

To test the importance of rapid glycolysis for actomyosin activity, we stained oxamate-treated and control amputated embryos for phosphorylated non-muscle myosin (pNM) and actin using immunohistochemistry and phalloidin, respectively. Phosphorylation of non-muscle myosin was unaffected by treatment with lower levels of oxamate, and only slightly reduced with higher concentrations (Fig. 4A,B). However, actin at the wound border was significantly diminished with both high and low concentrations of oxamate treatment. Myosin phosphorylation requires only a single ATP to donate the phosphate group for each myosin, and therefore is not the most energetically demanding process, whereas active contraction requires one molecule of ATP for each myosin stroke cycle (Spudich, 2001). We propose that it is this contraction that requires the use of glycolysis. It may be that actin stabilisation and condensation at the site of action requires activity of myosin, and the lack of actin remodelling may indicate a loss of active contraction.

**Figure 4.**
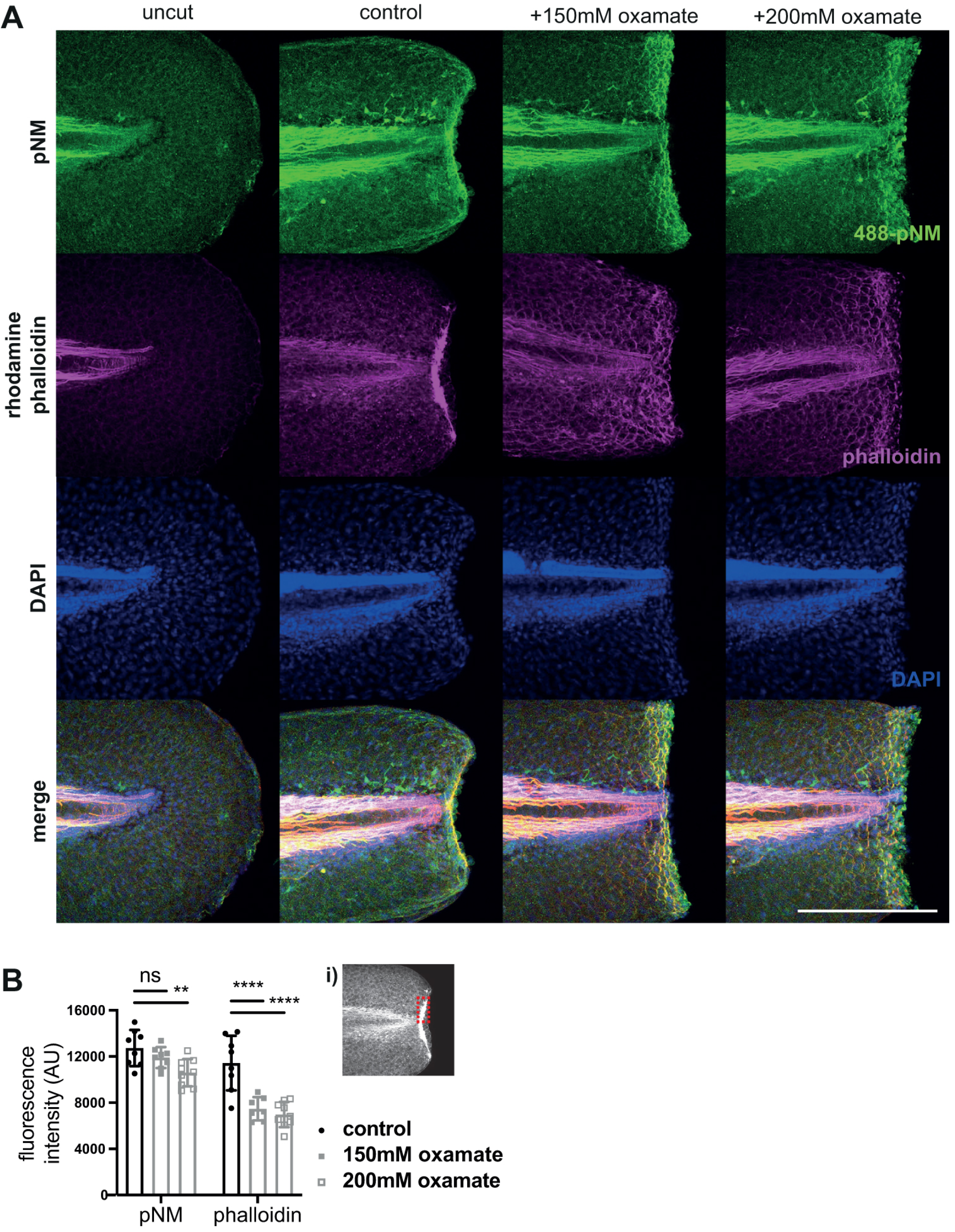
Effect of lactate dehydrogenase inhibition on actomyosin cable contraction in wound healing. (A) Maximal intensity confocal micrograph projections of representative embryos tails at 48hpf fixed and stained for phospho-non muscle myosin light chain II (pNM), actin, DAPI, and merged, at 10 minutes post amputation. (B) Graphs showing quantification of fluorescence intensity at the wound border of phalloidin actin staining and immunofluorescent pNM staining. Inset (i) denotes example of region measured. Two-way ANOVAs used to calculate significance, ** P<0.01, **** P<0.0001, ns P≥0.05, n=8. All scale bars represent 200μm.

### 3.4 Wound contraction is not dependent on oxidative phosphorylation

Our findings suggest that wound contraction is highly reliant on glycolysis, but this does not preclude a similar requirement for OXPHOS during wound contraction. Therefore, we endeavoured to determine whether inhibition of OXPHOS using NaN_3_ similarly affected wound contraction following larval tail fin amputation.

We performed fin fold amputations on 2dpf *Tg[ubb:laconic]*^*lkc1*^ embryos and immersed them immediately in NaN_3_ treatment, and then measured Laconic ratios after one hour of treatment. We found no significant difference in treated versus control embryos, although the NaN_3_ embryos trended toward having higher levels of lactate (Fig. 5A,B,C). Additionally, there was no difference in fin widths or wound contraction (Fig. 5A,C,D), thus we concluded that a reduction of mitochondrial activity does not appear to affect the rapid wound contraction phase following injury. However, given we did not measure wounds at a timepoint prior to 10mpa, when contraction has reached peak in control amputation, we cannot rule out the possibility that mitochondrial inhibition does not cause a brief acceleration in wound contraction.

**Figure 5.**
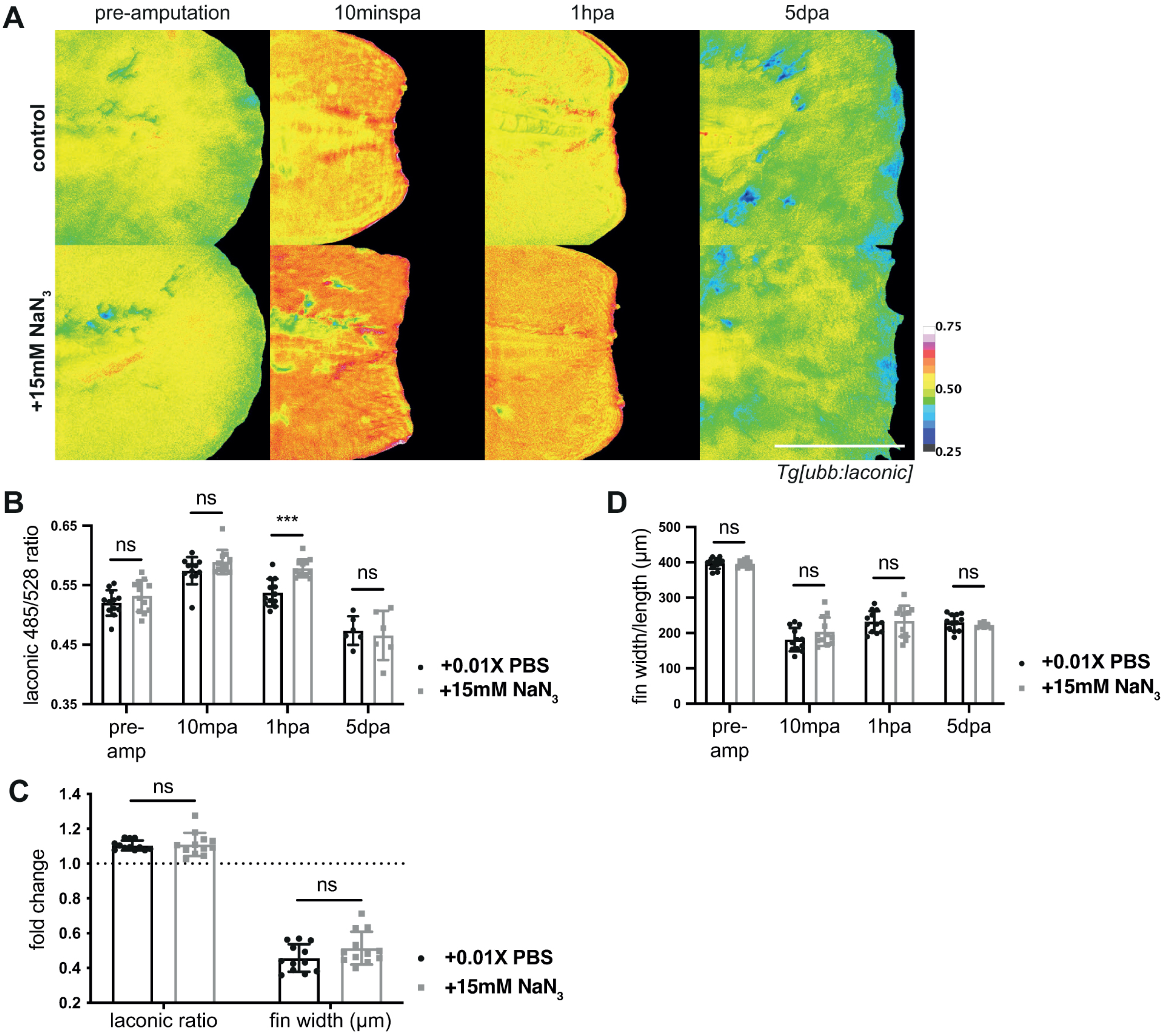
Mitochondrial inhibition in wound healing. (A) R Micrographs of representative *Tg[ubb:laconic]*^*lkc1*^ embryos tails treated at 48hpf for one hour post amputation with 15mM sodium azide (NaN_3_), to induce prolonged glycolysis and lactate production, or PBS control. Images were acquired at pre-amputation, 10 minutes, one hour, and five days post amputation, and pseudocoloured to show Laconic ratio. (B) Graph showing quantification of raw Laconic ratios pre-, 10 minutes post-, and five days post-amputation. Two-way ANOVA to calculate significance, n=12. (C) Graph showing fold change (10mpa value divided by pre-amputation value) of Laconic ratio and fin width in micrometres in the first 10 minutes of amputation with treatment with 15mM NaN_3_ or PBS control. Dotted line on the Y axis marks a fold change of 1 (no change). Two-way ANOVA to calculate significance, n=12. (D) Graph showing measured fin widths/length in micrometres pre-, 10 minutes post-, and five days post-amputation. Fin length measured from the tip of the notochord to the distal edge of the fin fold. Two-way ANOVA to calculate significance, n=12. All scale bars represent 200μm. Differences were considered significant to * P<0.05, ** P<0.01, *** P<0.001, **** P<0.0001, and ns P≥0.05.

By 1hpa control embryos have reduced in Laconic ratio, while NaN_3_ treated embryos continued producing elevated lactate levels (Fig. 5A,B). After washing out the drug at this point and allowing embryos to regenerate their fin folds, we imaged both conditions and found that there was no difference in Laconic ratio or fin regrowth at five days post amputation (dpa) (Fig. 5A,B,D). This suggests an overabundance of lactate in the early wound healing phase has no consequence on either wound closure or overall regeneration, while a reduction in lactate and glycolysis activity negatively impacts the rapid wound healing phase. Moreover, the apparent lack of requirement for OXPHOS in wound contraction suggests that this process is solely reliant on glycolysis.

### 3.5 Lactate is elevated in the notochord bead following tail amputations

After seeing that lactate levels rise dramatically, but only transiently, during the wound healing phase following amputation, we asked whether there was any evidence for metabolic reprogramming during the later regeneration phase, following either distal fin or tail amputation (Fig. S3A,B). During both larval fin fold and tail regeneration, a proliferative second phase occurs. However, following tail amputation an additional “notochord bead” arises from extruding notochord sheath cells, and displays high rates of proliferation (Romero et al., 2018). Overall, early lactate dynamics were similar following fin fold and tail amputations, with a near instant and rapid increase in Laconic ratio (Fig. 6A,B). However, the initial elevated lactate persists following tail amputations (24 hours post amputation (hpa)) when compared to distal fin fold amputations. In particular we found sustained higher Laconic ratios in the notochord bead until 48hpa (Fig. 6C). This blastema-like structure (Fig S3D) begins formation at 12 hours post amputation and blastema genetic markers begin to be lost after 48hpa (Kawakami et al., 2004; Romero et al., 2018); therefore, higher lactate levels correlated with the presence of the proliferative notochord bead and blastema-like structure.

**Figure 6.**
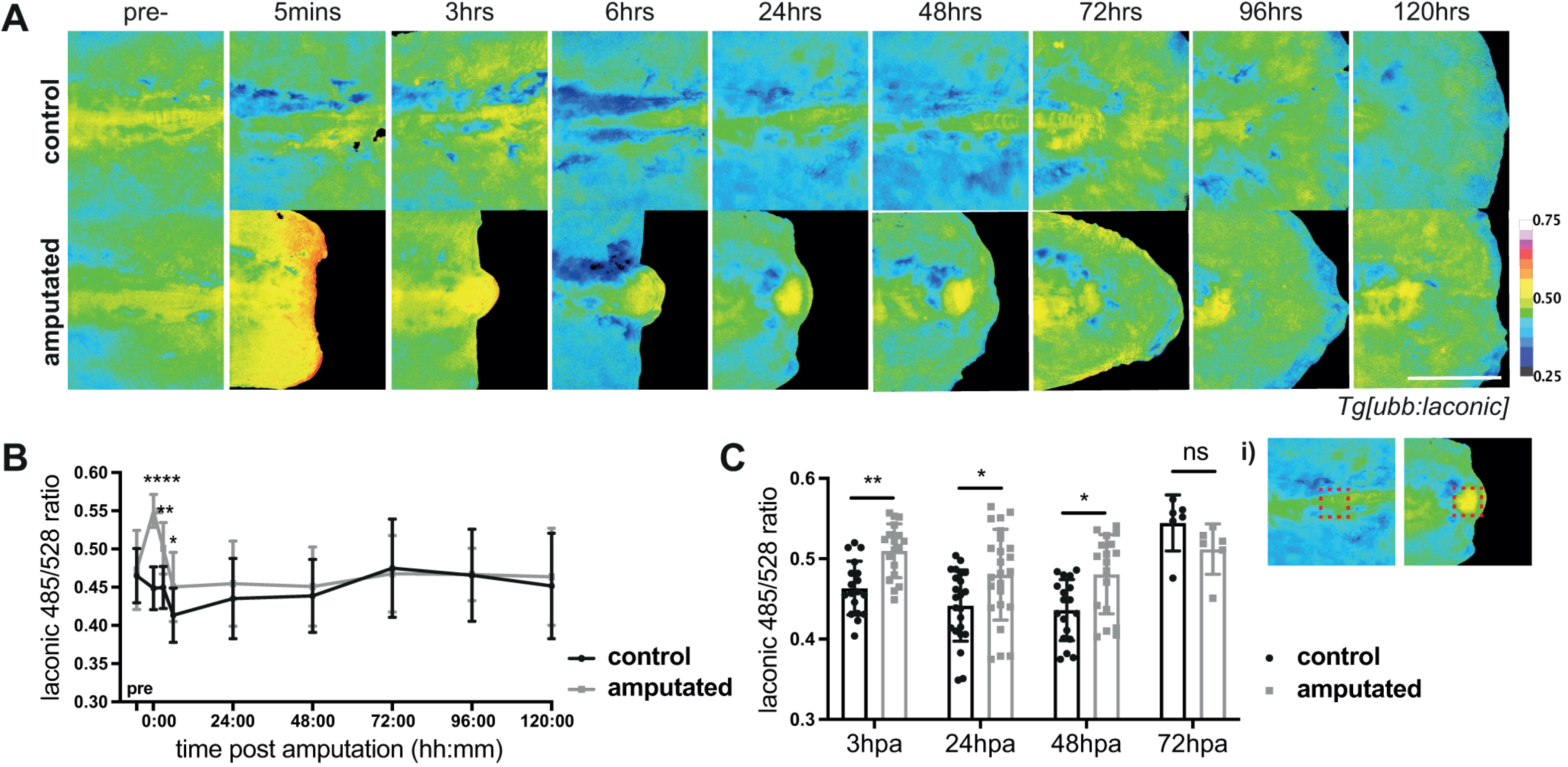
Lactate levels in tail regeneration. (A) Micrographs of representative *Tg[ubb:laconic]*^*lkc1*^ embryos tails at 48hpf imaged pre-amputation and at various time-points over the course of regeneration (note that images show representative embryos rather than following an individual embryo), pseudocoloured to show Laconic ratio. (B) Graphs showing quantification of raw Laconic ratios pre-treatment and at various timepoints post amputation. Two-way ANOVA to calculate significance, n=24. (C) Graph showing quantification of raw Laconic ratios specifically in the notochord bead region (as indicated in (i)). Students ‘t-test to calculate significance, n=24. Scale bar represents 200μm. Differences were considered significant to * P<0.05, ** P<0.01, *** P<0.001, **** P<0.0001, and ns P≥0.05.

### 3.6 Inhibition of glycolysis prevents successful tail regeneration

An up-regulation in glycolytic enzymes with an accompanying reduction in mitochondrial activity occurs in proliferating cardiomyocytes of regenerating zebrafish hearts (Honkoop et al., 2019) and genes involved in glycolysis and the PPP are significantly up-regulated during Xenopus tadpole tail regeneration (Love et al., 2014). Along with the elevation of aerobic glycolysis in the tail notochord bead noted previously, this prompted us to assess the potential role for aerobic glycolysis during zebrafish larval tail regeneration versus fin fold regeneration.

We thus treated 2dpf amputated *Tg[ubb:laconic]*^*lkc1*^ embryos over the course of regeneration with 2-deoxy-D-glucose (2DG), a competitive inhibitor of hexokinase, a critical enzyme in the glycolytic pathway (Wick et al., 1957) and analysed the resulting tail lengths at 5dpa. To confirm the efficacy of 2DG on lactate production, we measured lactate levels at 120hpa after treatment for the first 72hpa and verified that 2DG caused a significant decrease in Laconic ratio, and therefore lactate level, in both distal fin fold and tail amputation conditions at 120hpa and in 7dpf unamputated controls (Fig. 7A,B). Given that 2DG can potently reduce lactate levels in both our regeneration assays, we asked if regeneration of either the fin fold or tail was compromised in its presence. Tail amputated embryos did not regenerate when treated with 2DG, resulting in a significantly shorter tail length at 120hpa (Fig. 7C,D). In contrast, distal fin fold amputations, though averaging a shorter length of regrowth than controls, were not significantly affected (Fig. 7C,D), consistent with the lack of lactate increase in this post-wound healing phase. Later elevated Laconic ratios and therefore lactate levels were seen only in the notochord bead of tail amputations (Fig. 6A,C), while fin fold amputations lacked the formation of this structure and showed no significant difference in Laconic ratio throughout the duration of the regeneration phase (Fig. 2E). Thus, the importance of glycolysis is likely related to the formation of the blastema-like notochord bead structure. There was no difference between 2DG treated and control unamputated embryo fin lengths (Fig. 7C,D), suggesting glycolysis is not essential for normal embryo fin development.

**Figure 7.**
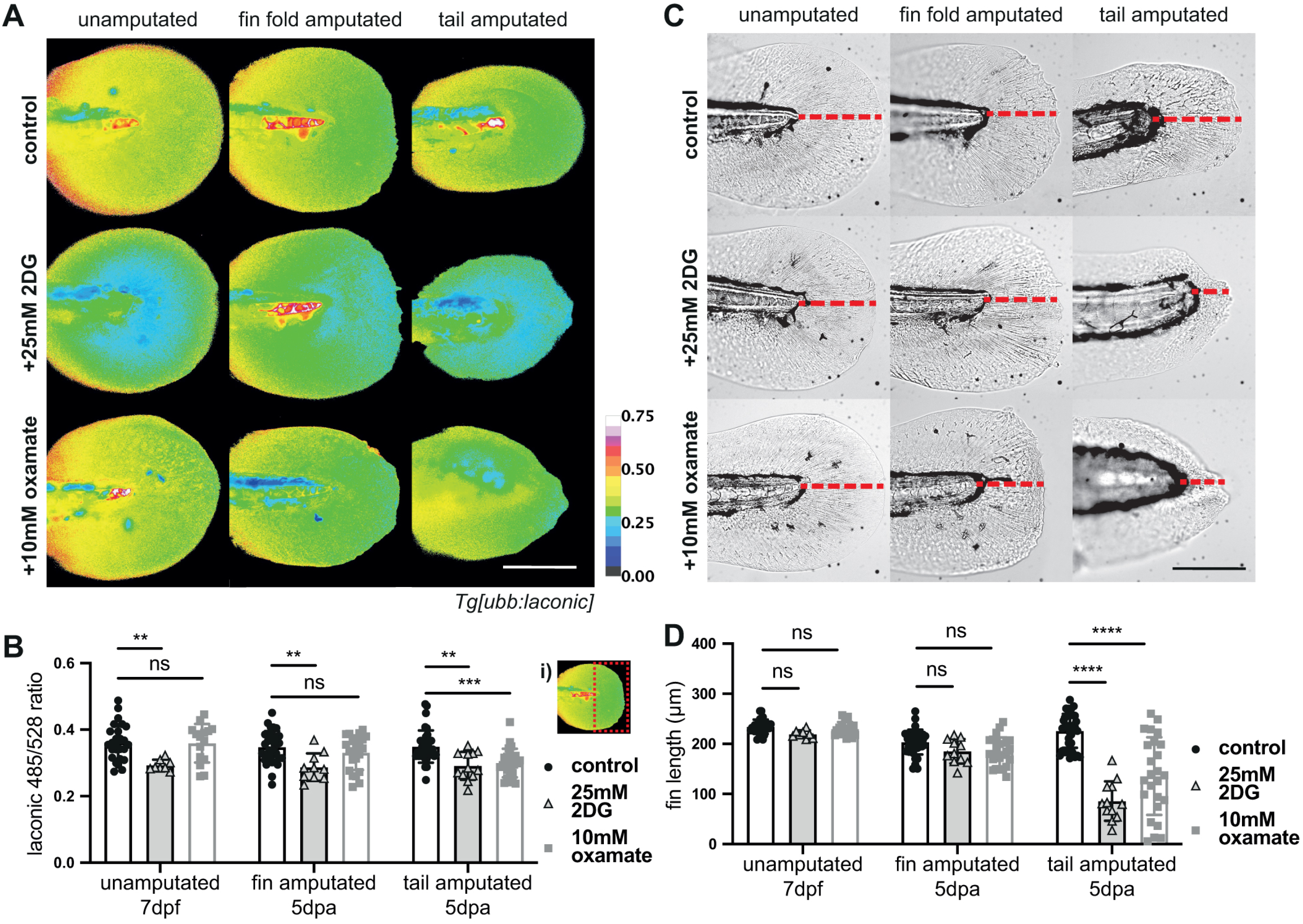
Glycolysis inhibition in regeneration. (A) R Micrographs of representative *Tg[ubb:laconic]*^*lkc1*^ larvae tails, pseudocoloured to show Laconic ratio. Imaged at 7dpf or 120hpa when amputated at 48hpf, treated for the first 72 hours post amputation with 25mM 2DG, for the full 120 hours of regeneration with 10mM oxamate, or with a control. (B) Graph showing quantification of raw Laconic ratios at 7dpf or 120hpa when amputated at 48hpf and treated for the first 72 hours post amputation with 25mM 2DG, 10mM oxamate, or control. Two-way ANOVA to calculate significance, n=13. White dashed box in inset (i) shows example of area measured for quantification of Laconic ratio in (B). (C) Brightfield images of representative *Tg[ubb:laconic]*^*lkc1*^ larvae tails at 7dpf or 5dpa when amputated at 48hpf, treated for the first 72 hours post amputation with 25mM 2DG, for the full 120 hours of regeneration with 10mM oxamate, or with a control. Red dashed line indicates measurement taken for fin length. (D) Graph showing fin length measurements taken at 7dpf or 120hpa when amputated at 48hpf and treated for the first 72 hours post amputation with 25mM 2DG or with a vehicle control. Two-way ANOVA to calculate significance, n=13. All scale bars represent 200μm. Differences were considered significant to * P<0.05, ** P<0.01, *** P<0.001, **** P<0.0001, and ns P ≥0.05.

To support the findings with 2DG, we additionally utilised oxamate, the LDH inhibitor we used previously to attenuate the initial burst of lactate production occurring during wound healing. The longer length of the regeneration experiments, compared to the wound healing assay, required a lower concentration of oxamate (10mM), as the higher concentrations used previously (150-200mM) were not compatible with survival over 5 days of exposure. Indeed, we noted that oxamate treatment visibly affected the swimming ability of the larvae as early as 24 hours after treatment. Nevertheless, we noted significantly reduced lactate levels in the tail amputated larvae treated with oxamate at 120hpa, although this reduction was not seen in fin fold amputated or unamputated larvae (Fig. 7A,B). As with the experiments with 2DG, oxamate treatment over the entirety of regeneration attenuated regrowth following tail amputation, while allowing distal fin fold amputated larvae to reach lengths comparable to those of the amputated controls (Fig. 7C,D). Likewise, oxamate treated embryos showed no difference in fin lengths when compared to unamputated controls (Fig. 7C,D) supporting the conclusion that rapid or aerobic glycolysis is not critical for normal fin formation of the embryos. Finally, to support the findings that attenuating LDH activity affected tail regeneration, but not distal fin regeneration, we also treated embryos with the alternative LDH inhibitor, GNE-140. As with oxamate, a lower concentration of the inhibitor was required (40mM) to sustain viability over the five days of treatment. Also like oxamate, GNE-140 treatment significantly reduced the regenerative ability of the larvae following tail amputation (Fig. S5A-B. Again, though in general regenerative length was shorter than controls, GNE-140 did not significantly affect regeneration following distal fin fold amputation.

Thus, we found that aerobic glycolysis plays an essential role during larval tail regeneration, which involves the regeneration of many tissues, including the spinal cord and notochord, but aerobic glycolysis is not required for distal fin fold regeneration, which is not associated with long-term elevation of lactate levels and is not dependent on a blastema-like structure.

## 4 DISCUSSION

### 4.1 The future of genetically encoded sensors *in vivo*

We have shown the genetically encoded sensor, Laconic, can successfully report lactate levels in zebrafish larvae. Use of genetic sensors of metabolites *in vivo* allow the assessment of the dynamic changes of metabolism with spatial resolution, a feat unachievable with biochemical methods. It further permits imaging with more refined temporal resolution, such as with time-lapse movies, and expressing the sensor under tissue specific promoters additionally enables selective interrogation of metabolites in distinct subtypes of cells during complex multicellular processes, such as organogenesis or regeneration.

Whilst effective, the Laconic sensor suffered from low fluorescence intensity in our transgenic line, which limited its sensitivity. There may also be additional interference from autofluorescence, especially in cells with high pigment or yolk content, and the multicellular nature of organisms, which additionally affects the sensitivity of the sensor. We were able to confirm the ability of Laconic to report lactate levels with a biochemical lactate assay, and, in future, combining genetically encoded metabolite sensors with biochemical assays, whole embryo metabolomics and/or MALDI mass spectrometry imaging will produce a complementary array of data, with the sensors providing a broader depiction of the temporal and spatial changes in metabolism while metabolomic approaches supplying a more comprehensive dataset of information of a large range of metabolites at given timepoints.

Other *in vivo* studies have also demonstrated the applicability of genetically encoded biosensors for measuring metabolite dynamics in zebrafish. The iNap1 sensor was used to show that NADPH levels decrease following embryonic fin amputation and co-localises with hydrogen peroxide (H_2_O_2_). This was interpreted as a result of dual oxidase (DUOX) activity, which consumes NADPH while generating H_2_O_2_ (Tao et al., 2017). Activity of the pentose phosphate pathway (PPP), specifically the enzyme glucose-6-phosphate dehydrogenase, is the main contributor to NADPH production (Spaans et al., 2015), and thus iNap sensors could in future studies be also utilised as an indicator of the Warburg effect alongside Laconic.

### 4.2 A role for aerobic glycolysis in early wound healing and formation of a blastema-like structure

In both distal larval fin fold and tail amputations the rapid increase in lactate levels within minutes following amputation occurs prior to the proliferative phase of regeneration, and correlates with the actomyosin contraction of the wound margin. Our chemical inhibitor experiments suggest that this rapid rise in lactate levels are necessary for wound contraction. More specifically, inhibition of LDH results in failure of the wound to contract and an attenuation of actin re-organisation and concentration at the wound border through purse-string action of myosin on actin. This reduction was not seen upon inhibition of mitochondrial OXPHOS activity, suggesting this process is solely dependent on glycolysis. One might ask whether this rapid rise in lactate levels immediately after amputation is the result of aerobic glycolysis or anaerobic glycolysis? Two lines of evidence point toward aerobic glycolysis. The first is that, as mentioned above, inhibition of OXPHOS has little effect on wound closure, thus rapid oxygen consumption due to OXPHOS is unlikely to be occurring. The second is that, based on evidence in *Xenopus* tadpoles, where there is a rapid rise in oxygen levels at the wound immediately following tadpole tail amputations (Ferreira et al., 2018), it is unlikely that the wound margin in our zebrafish embryos are becoming anoxic or hypoxic immediately after injury. A similar role for aerobic glycolysis has been shown in the brains of mice, whereby upon neuronal excitation increases glucose consumption, and glycolysis temporarily exceeds the rate of oxidative metabolism to provide for the rapid increase in energy demand (Díaz-García et al., 2017). Furthermore, the process of enucleation in erythrocytes requires contraction of an actomyosin ring and is prevented when aerobic glycolysis is blocked by inhibition of the glycolytic enzymes glyceraldehyde 3-phosphate dehydrogenase (GAPDH) or LDH (Goto et al., 2019). Thus, we propose that aerobic glycolysis provides a means of rapid ATP production necessary for driving the energy consuming process of actomyosin-mediated wound contraction minutes after amputation.

During the subsequent regeneration phases following wound healing, aerobic glycolysis, as indicated by lactate production, is once again implicated in the notochord bead/blastema in tail regeneration. In contrast, we saw no significant increase in lactate levels during fin fold regeneration. The larval fin fold “blastema” does not play a specific role in proliferation as it does in adult fin regeneration, and instead proliferation occurs in a more spatially distributed manner (Mateus et al., 2012). Tail amputation, however, is more similar to a canonical appendage regenerative response, in that multiple tissues must be replenished, including the notochord, spinal cord and skeletal muscle. The blastema formed following tail amputations also expresses *msx* genes and is partly made up of extruded notochord cells which create the “notochord bead” (Romero et al., 2018). We show that elevation of the Laconic ratio occurs as early as 3hpa and continues until 48hpa and is restricted to the notochord bead. The raised lactate levels correlate temporally with the blastema, returning to control levels after 48hpa as the regenerant enters the third phase of regeneration, characterised by differentiation and progressive scaling back of proliferation (Kawakami et al., 2004; Mateus et al., 2012). Other work has also shown elevated levels of glycolysis gene expression and decreased mitochondrial activity during zebrafish heart regeneration. Inhibition of glycolysis with 2DG resulted in a reduction of proliferating cardiomyocytes (Honkoop et al., 2019), indicating this metabolic switch to glycolysis is required for regrowth. Increased expression of glycolysis genes has also been observed in zebrafish following larval tail amputation, with glycolysis inhibition resulting in abnormal blastema formation (Sinclair et a., 2021), and we additionally find that activity of the glycolytic enzymes hexokinase and LDH are required for larval tail regeneration. Thus, aerobic glycolysis is required for successful regeneration through the formation or output of the blastema-like notochord bead. The regrowth of the epidermis following fin fold amputations, however, does not require the function of these glycolytic enzymes and achieved regrowth comparable to controls despite glycolytic inhibition. It is unclear at this point if this different reliance on aerobic glycolysis between the two amputation models reflects diversity in the constituent cell types being regenerated or the differing anabolic needs for regeneration of the fin fold, versus overall regeneration of many tissue types. Intriguingly, recent findings suggest that a similar metabolic switch also occurs following adult fin regeneration in zebrafish, and inhibition of this switch results in failure in blastema formation in the adult fin as well (Brandão et al., 2022).

Thus, our work suggests that aerobic glycolysis is important at two distinct points following injury; the first being within minutes during the rapid wound healing phase and the second during the tail regeneration phase. Though a blastema is typically highly proliferative, there is an absence of raised lactate levels in any region aside from the notochord bead during the proliferative phase of fin and tail regeneration. Aside from being a product of the Warburg effect, lactate may also have a direct effect on blastema formation and function, such as acting as a second messenger. For example, lactate has recently been shown to mediate magnesium uptake into the mitochondria (Chubanov & Gudermann, 2020), which in turn has been reported to have a stimulatory effect on oxidative metabolism and may affect mitochondrial calcium flux (reviewed in Pilchova et al., 2017). The downstream targets and signalling stimulated by lactate in this instance remain unknown, their elucidation a possible direction for future studies. Other future work could also look into whether proliferative cells are reduced in glycolysis-inhibited tail amputations, as is the case in zebrafish heart regeneration (Honkoop et al., 2019).

The underlying mechanisms governing metabolic reprogramming during tail regeneration remain unknown. Both hypoxia inducible factor-1α (HIF1α) signalling and the embryonic form of pyruvate kinase (PKM2) have been implicated in the switch of induced pluripotent stem cells to glycolytic metabolism, leading to their de-differentiation (Prigione & Adjaye, 2010; Prigione et al., 2014). More broadly, there is increasing evidence that hypoxic conditions and reactive oxygen species (ROS) influence glycolytic switching. HIF1α signalling is also sufficient for inducing reprogramming to glycolytic metabolism in mouse embryonic stem cells (W. Zhou et al., 2012) and is known to have a positive effect on glycolysis, such as in cancer (reviewed in Nagao et al., 2019) and macrophages (T. Wang et al., 2017). Further, H_2_O_2_ has been shown to positively regulate glycolysis in cancer cells (Shi et al., 2009). Illuminating the molecular pathways involved in successful regeneration, such as the relationship between H_2_O_2_ and glycolysis, will assist in determining the logic of metabolic reprogramming in different phases of regeneration. Zebrafish imaging approaches, combining an expanding genetically encoded biosensor toolbox with high regenerative capacity, offers a unique system to determine principles of metabolic programming in regeneration.

## ACKNOWLEDGEMENTS

We would like to thank Peter March from the core imaging facility at UoM for help with imaging and the aquarium staff in the BSF unit (University of Manchester), and IMCB and LKCMedicine zebrafish facilities for their care of the fish.

## AUTHOR CONTRIBUTIONS

Conceptualisation: C.A.S., T.J.C, E.A.; Methodology: C.A.S.; Validation: C.A.S.; Formal analysis: C.A.S.; Investigation: C.A.S.; Writing - original draft: C.A.S.; Writing - review & editing: C.A.S., T.J.C, E.A.; Supervision: T.J.C, E.A.; Project administration: E.A.; Funding acquisition: E.A.

## SUPPLEMENTARY FIGURE LEGENDS

**Supplementary Figure 1.**
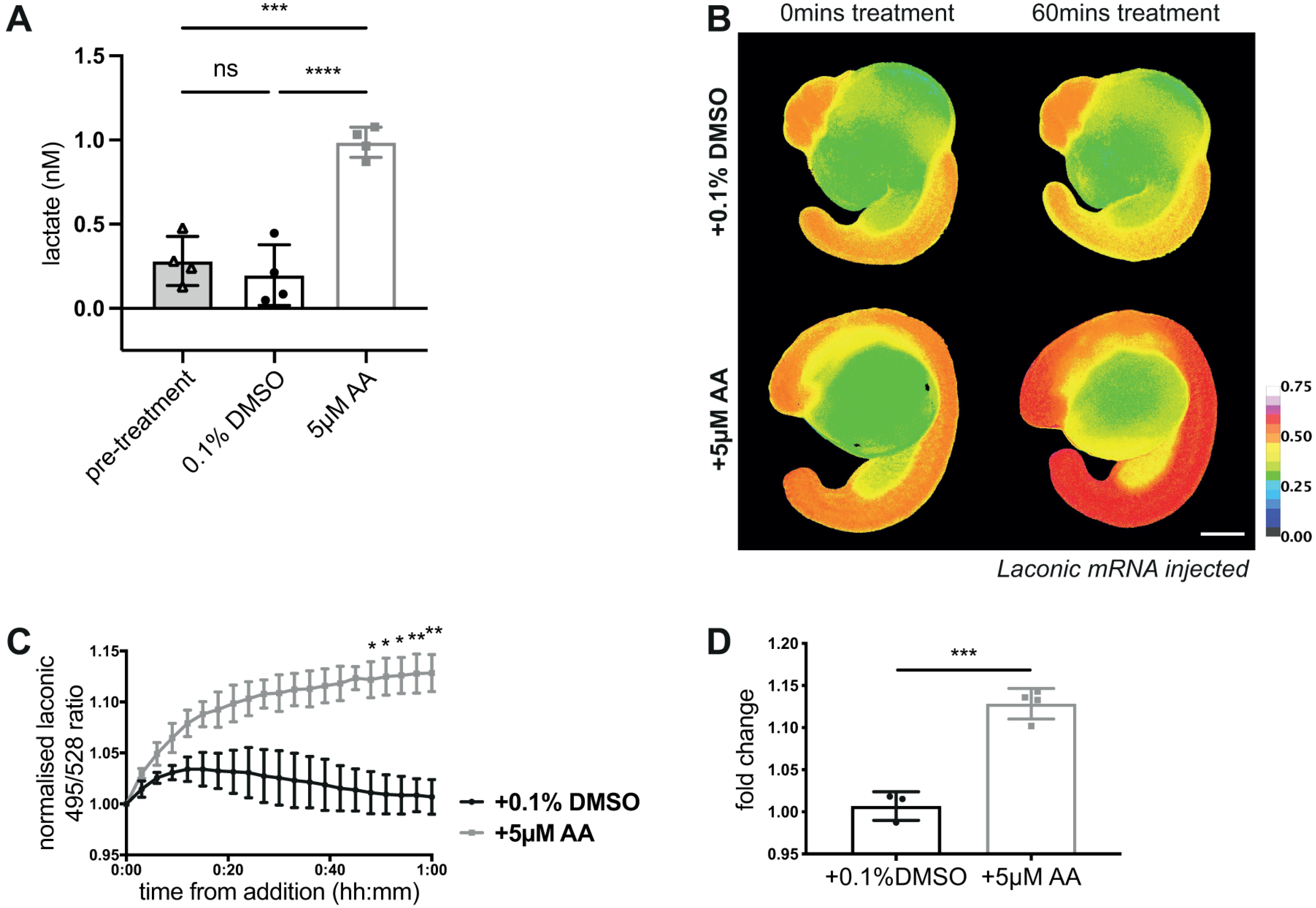
Schematic diagrams of the fin and tail amputation models of zebrafish embryo regeneration. (A) Schematic drawing of a two-day post fertilisation zebrafish embryo. e: eye, ov: otic vesicle, h: heart, y: yolk, s: somites, tf: tail fin. Red dashed box indicates the area shown in panel (B). (B) Tail region with amputation planes for fin fold and tail amputations (dashed red lines). A fin fold amputation site is positioned just distal to the tip of the notochord and excision is limited epithelial tissue and mesenchymal cells, while a tail amputation is oriented using the pigment gap and circulatory loop of the caudal vein, severing notochord, neural tube, and muscle in addition to epithelial tissue. s: somites (yellow), n: notochord (pink), cv: caudal vein (blue), p: pigment (black), tf: tail fin fold. (C) General timeline for amputation experiments, with phases of regeneration and inhibitor treatment times used in Figure 7 indicated. Embryos are imaged at various time points depending on the specific experiment between ten minutes post amputation (10mpa) and full regeneration at five days post amputation (5dpa). Divisions are 24 hours. (D) Schematic depiction of the general timeline for regeneration. Embryos are amputated at 48hpf, following which the wound contracts and closes within ten minutes. Over the next five days the embryo regenerates the removed tissue, divided into three phases. The events are equivalent in both fin fold and tail amputations.

**Supplementary Figure 2.**
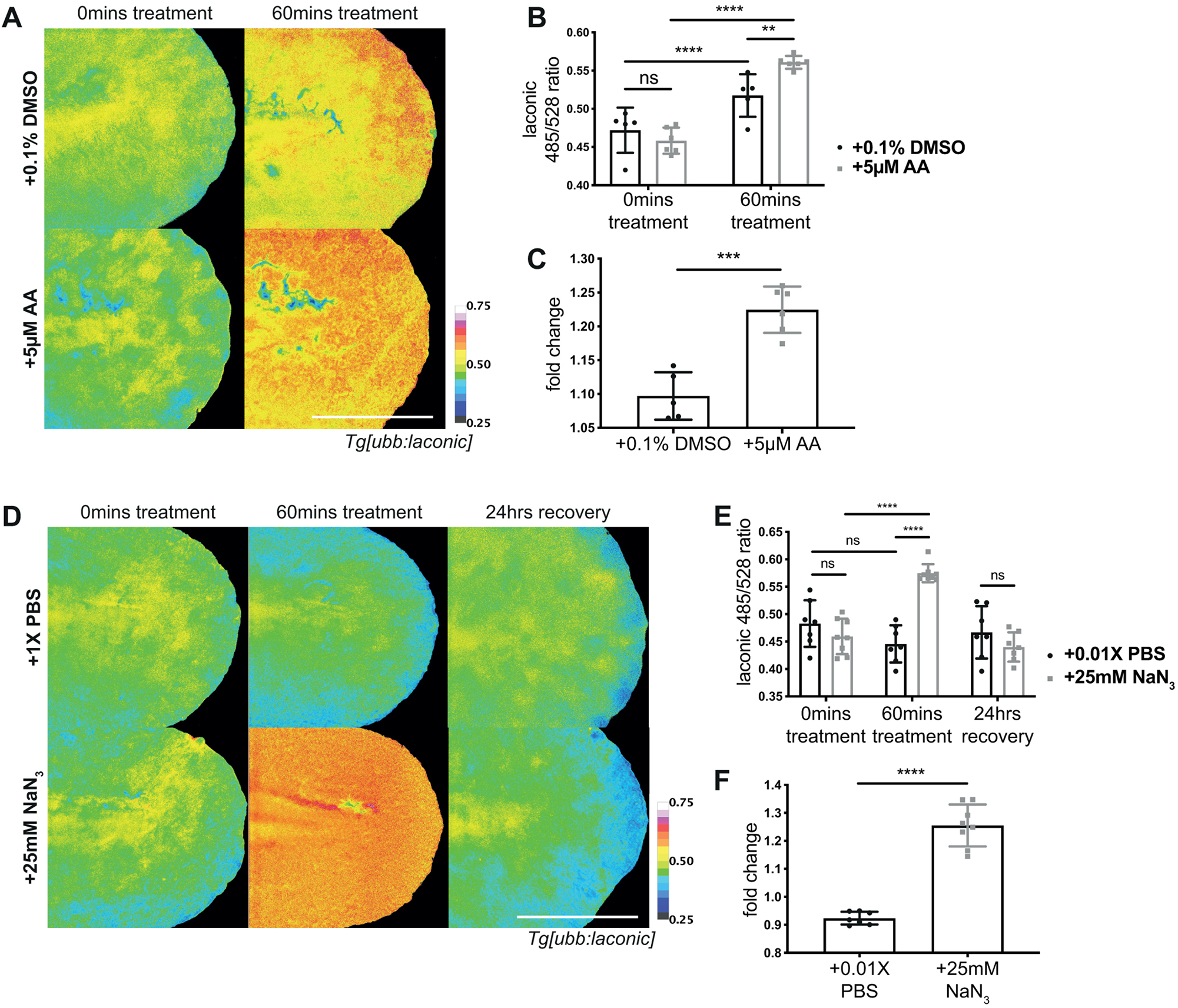
Laconic positive controls as mRNA injections. (A) Graph of lactate levels in 2dpf embryos (calculated using a standard curve) after 10 minutes treatment with 0.1% DMSO or 5μM AA compared to a pre-treatment baseline level. One-way ANOVA to calculate significance, n=36. (B) Micrographs of representative ∼19hpf wild type embryos injected with laconic mRNA at the one cell stage before and after 60 minutes of treatment with 0.1% DMSO or 5μM AA, pseudocoloured to show Laconic ratio. (C) Graph showing Laconic ratios over time during treatment with 0.1% DMSO or 5μM AA, normalised to pre-treatment value. Two-way ANOVA to calculate significance, n=4. (D) Graph showing quantification of ratio change as fold change (post-treatment ratio divided by pre-treatment ratio). Students ‘t-test to calculate significance, n=4. All scale bars represent 200μm. Differences were considered significant to * P<0.05, ** P<0.01, *** P<0.001, **** P<0.0001, and ns P≥0.05.

**Supplementary Figure 3.**
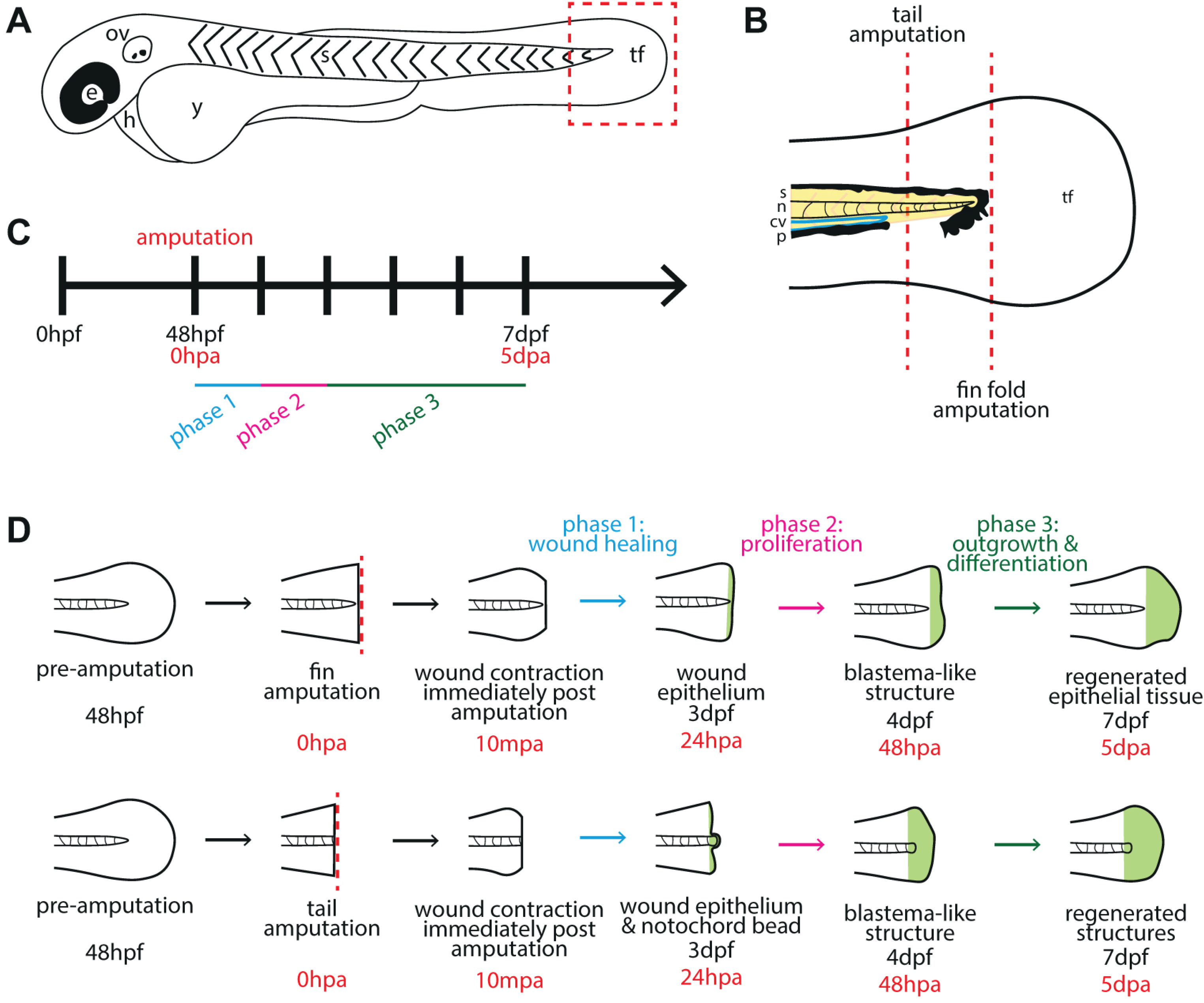
Laconic positive controls as a transgenic line. (A) Micrographs of representative *Tg[ubb:laconic]*^*lkc1*^ embryos tails at 48hpf before and after treatment with 0.1% DMSO or 5μM AA, pseudocoloured to show Laconic ratio. (B) Graph showing raw Laconic ratios pre-treatment and after treatment with 0.1% DMSO or 5μM AA. Two-way ANOVA to calculate significance, n=6. (C) Graphs showing quantification of ratio change after 60 minutes of treatment with 0.1% DMSO or 5μM AA as fold change (ratio after treatment divided by pre-treatment value). Students ‘t-test to calculate significance, n=6. (D) Micrographs of representative *Tg[ubb:laconic]*^*lkc1*^ embryos tails at 48hpf before and after 60 minutes of treatment with 0.01X PBS or 25mM sodium azide (NaN_3_), and after 24 hours recovery post wash out of drug, pseudocoloured to show Laconic ratio. (E) Graph showing raw Laconic ratios pre-treatment, after 60 minutes of treatment with 0.01X PBS or 25mM NaN_3_, and after 24 hours of recovery after drug wash out. Two-way ANOVA used to calculate significance, n=8. (F) Graph showing quantification of ratio change after 60 minutes of treatment woth 0.01X PBS or 25mM NaN_3_ as fold change (ratio after treatment divided by pre-treatment value). Students ‘t-tests used to calculate significance, n=8. All scale bars represent 200μm. Differences were considered significant to * P<0.05, ** P<0.01, *** P<0.001, **** P<0.0001, and ns P≥0.05.

**Supplementary Figure 4.**
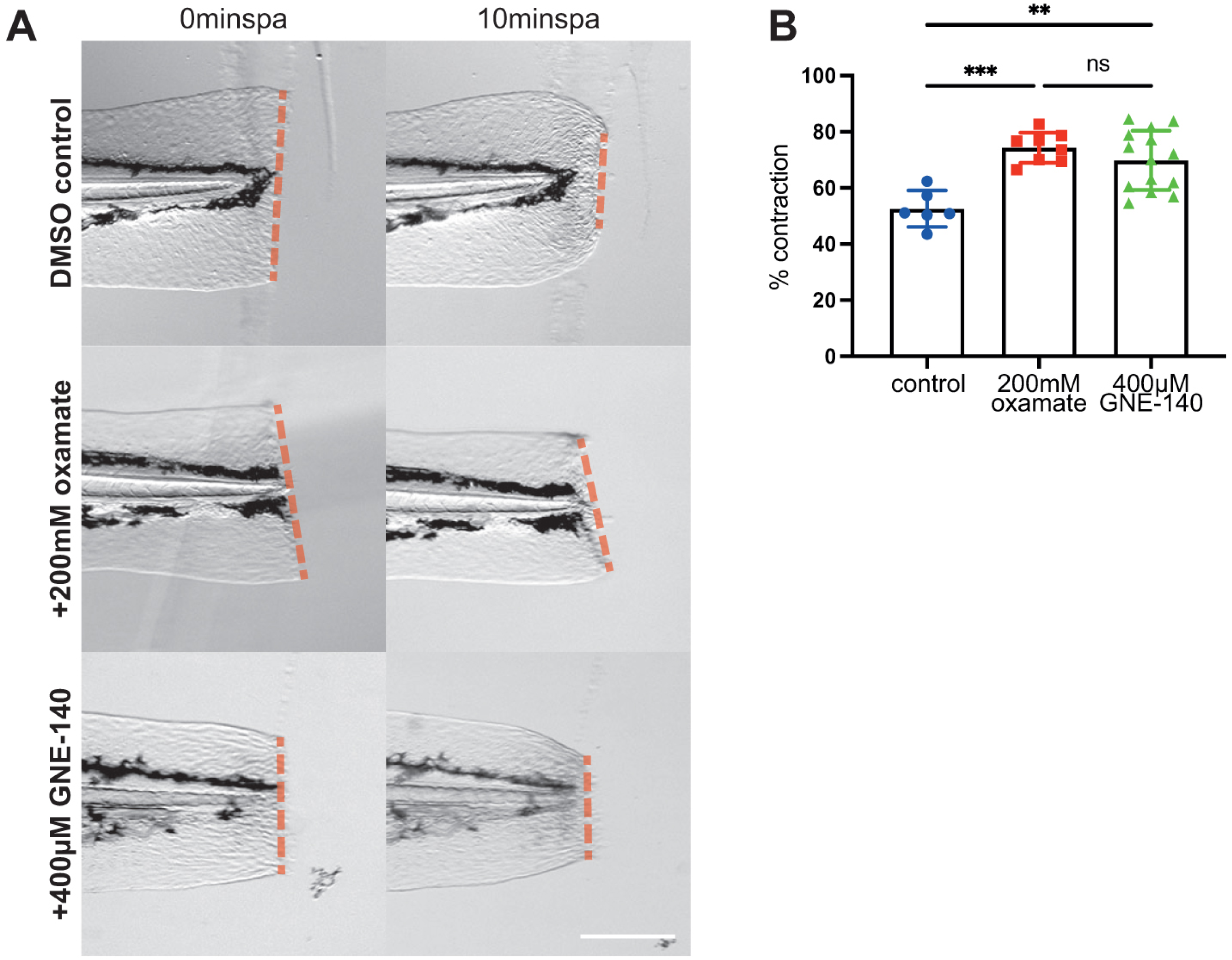
Further inhibition of lactate production during fin fold amputation. (A) Brightfield images of 2dpf WT control (no treatment) compared with 200mM oxamate- and 400µM GNE-140-treated embryos at 0minspa and 20minspa. Red dashed lines indicate example measurements taken for quantification. Scale bar represents 200µm. (B) Graph showing percentage contraction at 10minspa of wound width at 0minspa. One-way ANOVA to calculate significance.

**Supplementary Figure 5.**
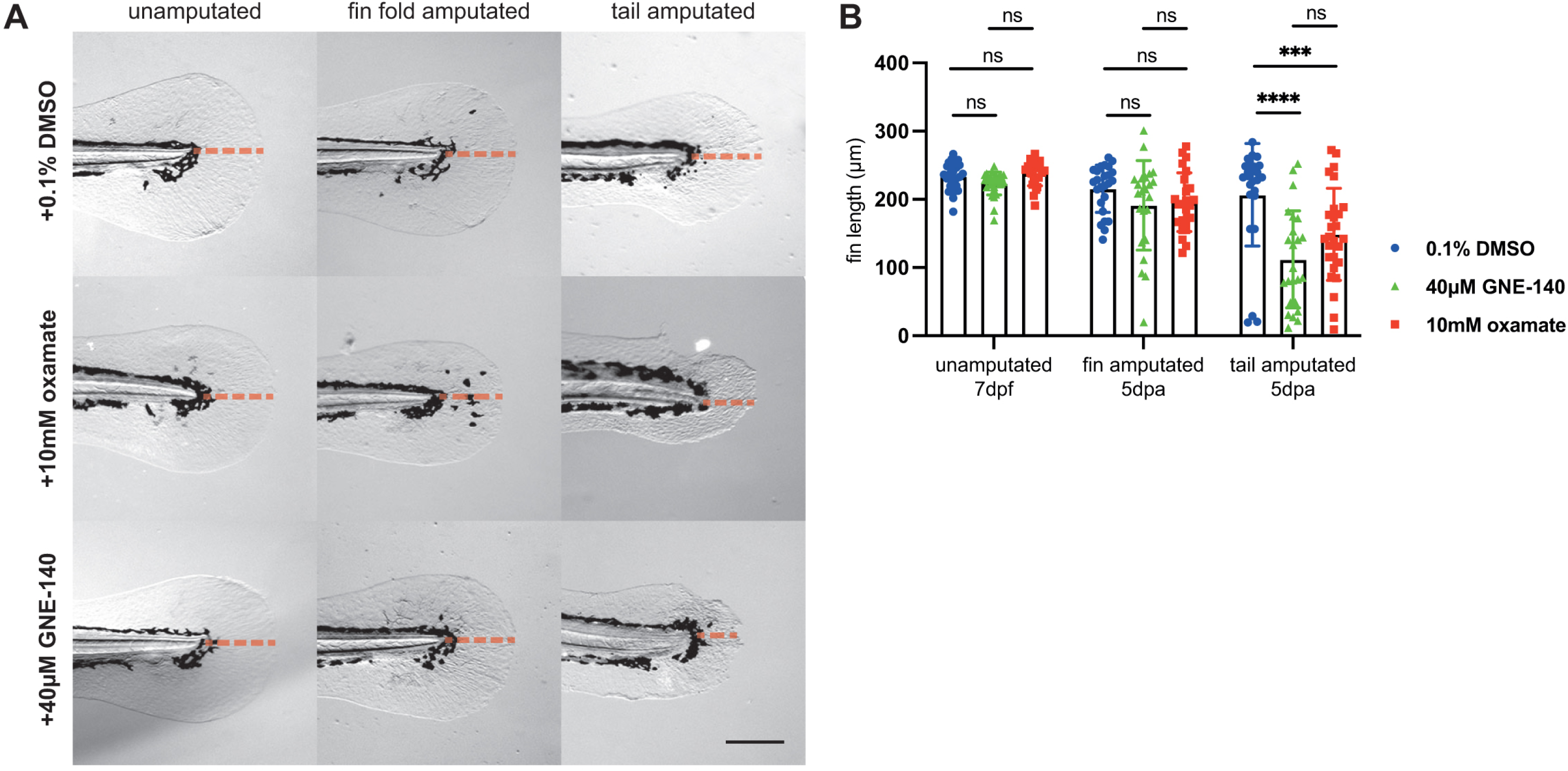
Further inhibition of lactate production over the whole of regeneration. (A) Brightfield images of representative WT embryos at 5dpa (7dpf), treated with 0.1% DMSO (control), 10mM oxamate, or 40µM GNE-140. Red dashed line indicates measurements taken for quantification of regrowth. Scale bar represents 200µm. (B) Graph comparing inhibited (10mM oxamate or 40µM GNE-140 treatment) with control (0.1% DMSO treatment) embryos in the unamputated, fin fold amputated, and tail amputated conditions at 5dpa (7dpf). Two-way ANOVA to calculate significance.

## SUPPLEMENTARY MOVIE LEGENDS

Supplementary Movie 1. Time-lapse movie generated of the rate of wound healing / contraction following distal fin amputation under control / untreated condition over 12 minutes. 360 still images were taken every 2 seconds over 12 minutes and rendered as a time-lapse movie using iMovie at 40 frames per second, giving a 9 second time-lapse movie. https://youtu.be/3cqNSHLE0iA

Supplementary Movie 2. Time-lapse movie generated of the rate of wound healing / contraction following distal fin amputation under 200mM oxamate treated condition over 12 minutes. 360 still images were taken every 2 seconds over 12 minutes and rendered as a time-lapse movie using iMovie at 40 frames per second, giving a 9 second time-lapse movie. https://youtu.be/XFSCztl_ep0

Supplementary Movie 3. Time-lapse movie generated of the rate of wound healing / contraction following distal fin amputation under 1% DMSO control treated condition over 12 minutes. 360 still images were taken every 2 seconds over 12 minutes and rendered as a time-lapse movie using iMovie at 40 frames per second, giving a 9 second time-lapse movie. https://youtu.be/jPAHR3AG6Og

Supplementary Movie 4. Time-lapse movie generated of the rate of wound healing / contraction following distal fin amputation under 400µM GNE-140 treated condition over 12 minutes. 360 still images were taken every 2 seconds over 12 minutes and rendered as a time-lapse movie using iMovie at 40 frames per second, giving a 9 second time-lapse movie. https://youtu.be/jIdOq-6TFDw

**Supplementary Table 1.**

| Insert | Original backbone | New backbone | Restriction enzyme 5' | Restriction enzyme 3' | Double digest conditions | New construct | Notes |
| --- | --- | --- | --- | --- | --- | --- | --- |
| Laconic | pcDNA3.1(-) | pCS2+ | EcoRI | XbaI (site added from HindIII site via short oligos) | Roche buffer H, 37°C | pCS2-Laconic | 1. Digest pcDNA3.1-Laconic with HindIII (Roche buffer B, 37°C), XbaI site added by ligation with primers*<br>2. Digest new pcDNA3.1-Laconic with EcoRI and XbaI (2262bp)<br>3. Digest pCS2 with EcoRI and XbaI (4074bp) (will create extra 40bp fragment)<br>* 5'-AGCTG<u>TCTAG</u>ACCCA-3'<br>5'-AGCTTGGG<u>TCTAG</u>AC-3' |
| Laconic | pcDNA3.1(-) | p3-HyPer | BamHI | HindIII | Roche buffer B, 37°C | p3-Laconic | 1. Digest Laconic with BamHI and HindIII (2239bp)<br>2. Digest p3-HyPer with BamHI and HindIII (2967bp) |

Laconic/pcDNA3.1(-) was a gift from Luis Felipe Barros (Addgene plasmid #44238; http://n2t.net/addgene:44238; RRID:Addgene\_44238) (San Martin et al., 2013).

